# Alternative mating tactics and tactical deception: Is the common cuttlefish (*Sepia officinalis*) nothing but a common cheat?

**DOI:** 10.1101/226191

**Authors:** Gavan M Cooke, Claire Johnson, Tony Reed, Andrew C Jackson

## Abstract

Alternative mating tactics (AMTs) are common in the animal kingdom, yet much work remains before their evolution and role in driving sexual selection is fully understood. Utilizing features of citizen science, we present compelling evidence that a third species in the cuttlefish genus *Sepia* (Cephalopoda) possess males who use sneaky mating and female mimicry as alternative strategies to conspicuous signalling and fighting. We also present new evidence of large aggregations (n~30) in this species that possibly drive alternative mating strategies. Lastly, we provide footage of an opportunistic sneaky copulation in this species. We believe that alternative mating tactics may be more common in this genus than previously recorded (based on observations presented here and a search of the literature for similar life history, environmental and behavioural factors found in other species within the *Sepia* genus), and as much of their captive husbandry is well known, they could an ideal system for studying the evolution of alternative reproductive strategies.

## 1.0 Introduction

Alternative mating tactics (AMTs) are common in the animal kingdom (Gross, 1996; Neff and Svensson 2013) and are seen in many different taxa (e.g. sunfish (*Lepomis macrochirus);* ruffs (*Philomachus pugnax*); swordtail (*Xiphophorus* sp.); side blotched lizard (*Uta stansburiana*) see table 1 Neff and Svensson 2013). A mating tactic can be fixed genetically (e.g. damselflies (*Ischnura* spp.) Cordero, 1990) or be conditional based (e.g. a dung beetle (*Onthophagus* sp.) Emlen 1994) where size at maturity determines the strategy. Different AMTs have evolved, such as; resource guarding (mates and food/refuges – the most commonly seen strategy and to which other tactics are normally alternative); female mimics (to reduce competition by deceiving other competitors); sneakers (to mate with a female whilst a competitor is distracted but would otherwise try and prevent) and examples of all three have evolved in a marine isopod (*Paracerceis sculpta*; Schuster 1991) for example. Males are frequently larger and more conspicuous than females, and this is likely a result of the necessity for intrasexual competition due to asymmetrical investment in reproduction between the sexes (Andersson, 1994), a feature seen widely throughout the animal kingdom (birds, Zi et al., 2003; fish, Endler, 1983; mammals and insects, Gage et al., 2002). Asymmetries arise due to the differences between the cost of producing sperm versus eggs, and commonly plays out in females being choosy of males and males competing over females. Although females may use AMTs (Neff and Svensson, 2013), males are more likely to adopt an alternative tactics as result this competition for females. If males are to adopt AMTs, it is predicted that smaller males are more likely too, compared to larger males, who are more likely to benefit from using honest, conspicuous sexually selected signals that possess a cost (Gross, 1996; Reynolds, 1996; Taborsky 1998).

**Table 1.** Ethogram of the behaviour described as deception from supplementary video Movie S1

| Behavioural element | Description |
| --- | --- |
| Mantle flare | Right side of mantle-head opening opens wide |
| Arm hide | Arms R3 and R4 are curled under and then within the newly created opening in the mantle |
| Body leaning | The opposite side of the cuttlefish leans towards the female |
| Alternative arm presentation | The unhidden L4 arm moves towards the female |
| Arm blow out | Cuttlefish appears to forcibly 'blow' the hidden arms from out of the mantle cavity which closes |

Males are often most conspicuous during the acquisition of mates because during this time they may use elaborate displays and signals that are meant to attract the attention of females. Conspicuous signals often possess significant costs (Zhavi, 1975); the costs ensure the signal presented represents the signaller’s qualities in some way, and have become known as ‘honest signals’ (see Zhavi, 1975). Although natural selection should favour honest signals (Zhavi, 1975), deception may be seen at low frequencies (Johnstone and Grafen 1993) if honest signals succeed most of the time. If a male cannot pay the cost of an elaborate signal (perhaps due to reduced foraging opportunities or being unable to avoid predation successfully) the male may have to change tactics and, indeed, AMTs are positively associated with exaggerated secondary sexual characteristics used as signals in courtship (Neff and Svensson 2013).

In addition to the potential costs of being seen by a predator, or the time taken to obtain the correct essential foods for an honest signal, animals may inadvertently signal to more animals than they intend to. Well established theory in animal communication suggests animal signals are a form of exploitation by the signaller of the intended receiver (Guilford and Dawkins, 1991), however, a signal may also be received by eavesdroppers who may be able to gain information about the signaller (McGregor, 1993) and their intentions (e.g. court/copulate with a female). A signaller may try and reduce the cost of eavesdropping through tactical deception (Kuczai et al. 2001) i.e. signals may not consistently reveal the genuine status or intentions of the signaller. To be a genuine a case of deception, the receiver of the deception must pay a cost (e.g. Kuczai et al. 2001) and the deceiver must benefit (Byrne and Whitton 1992; Hauser 1997).

Another important influence on AMTs is the operational sex ratio (OSR: the ratio of sexually mature males to fertilisable females - Mills and Reynolds, 2003) within a breeding population. If the OSR is skewed towards more males, then this increased competition for females can further drive sexual selection, including AMTs. OSRs are variable between species and even within them from population to population. This variation in the distribution of sexually mature adults may arise from environmental influences on the potential reproductive rates of the sexes (Kvarnemo 1994; Reynolds et al. 1986).

## 1.1 AMTs and cuttlefish

Brown et al. (2012) reported an AMT using tactical deception in *Sepia plangon* (a cuttlefish of the Class Cephalopoda) where males use a ‘split’ signal. The split signal includes male mating signals (zebra patterning; see Hanlon and Messenger, 1996) on one side of the body (the side towards a female) and a female pattern (e.g. blotched) on the other side – which is directed towards a rival male. In certain circumstances, this could allow males to mate with females without alerting other males i.e. a sneak AMT. The authors suggest this strategy deceives a conspecific whilst simultaneously attracting a female (see figure 1, Brown et al. 2012). Alternatively, two further studies (Norman et al. 1999; Hanlon et al. 2005) describe a different AMT in another cuttlefish (*Sepia apama*) where small males adopt female colouration and postures to gain access to females, even while dominant mate guarding males are present (female mimicry based AMT).

**Fig 1.**
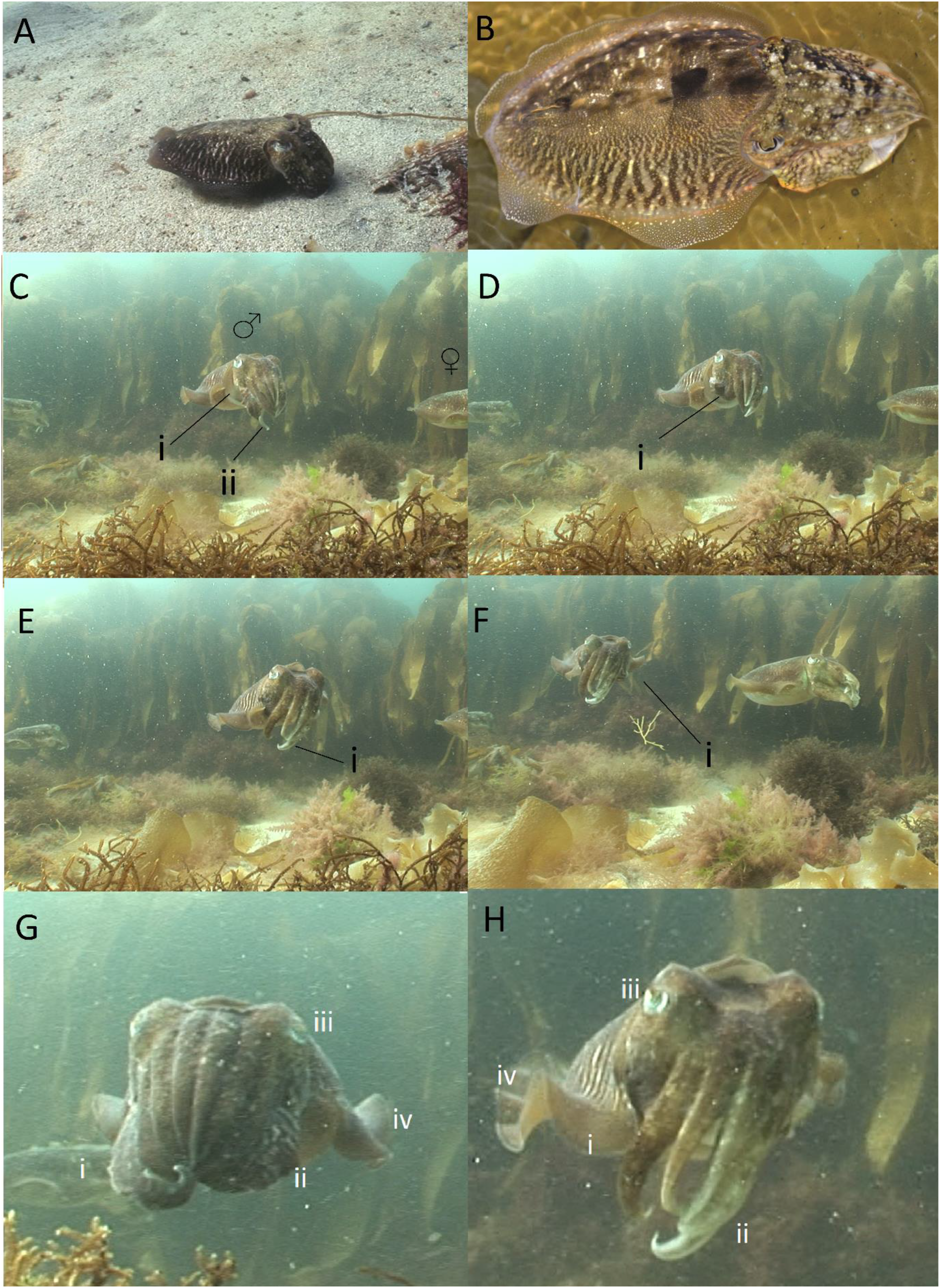
A) Still from a video (Video 1, supplementary material) taken of a cuttlefish employing the split signal display by a *S. officinalis* male. B) photograph from an aquarium raised cuttlefish from 2014. A male was observed to the left of this cuttlefish (out of shot) and a female to the right of (also out of shot). Note: zebra patterning does not have the same intensity as when presented to a rival male (Figure 2C/E). C-F) Sequence of events from supplementary video Movie S1 showing the sneaky behaviour by a male. C) Male (M) flares open the mantle opening (i) on its right-hand side whilst next to a likely female. D) Arms R3 and R4 (Cii and Di) curl under the remaining arms and into the gap created. E) Male begins to lean into the female and highlights his left-hand side, waving the unhidden male characteristics on its lower L4 arm (F). (G/H) There are clear differences in the postures and behaviours before the arm tucking and during, such as absence of R3 and R4 (i); the uncurling and brightness of arms L3 and L4 (ii); alertness as exhibited by wider eyes (iii) and increased mantle fin oscillations (iv). Stills A, C-H taken from video recorded in September 2013 by A. Jackson. Photograph (B) taken by B. Tonkins.

Although better known for their near whole body real time adaptive camouflage (e.g. Hanlon and Messenger, 1998), cuttlefish possess transient secondary sexually characteristic signals e.g. the intense zebra patterning used by the common cuttlefish (*Sepia officinalis,* Hanlon and Messenger 1998). This trait, whilst used in male-male competition as a signal of aggressiveness, which may settle contests before physical contact is required, is apparently repellent to females who may wish avoid aggression (Boal 1997). Small patches of epidermis that are less chromatically changeable are also found on mature males (Hanlon and Messenger 1998). These are bright, reflective patches of pigments (leucophores/iridophores) on the male’s lower outer arms (Hanlon and Messenger, 1998), *S. apama* hide theses outer arms when Adopting a female mimic strategy (Norman et al. 1999; Hanlon et al., 2005). *S. officinalis* is a well-known and widely studied cuttlefish species in a laboratory setting (Smith et al. 2013) but still little is known about their behavioural ecology in natural settings (Adamo et al. 2000). Fortunately, several fisheries based studies (e.g. Boletzky, 1983; Dunn 1999; Bloor et al. 2013a; Gauvit et al. 1997) are available for the region of the cuttlefish observed here (south Cornwall, UK) which provide invaluable data. They are sexually dimorphic for size (larger males (Dunn 1999), but this can be subtle and difficult to when individuals are alone or not fully mature) and both sexes exhibit seasonal migrations (Boletzky, 1983; Dunn 1999; Bloor et al. 2013a; Gauvit et al. 1997), leaving the shallow waters where they hatched to overwinter at depths <200m, returning the next spring. The English Channel is thought to have a biennial population (Bloor et al, 2013a) that is assumed to be semelparous, as are all cuttlefish (Hanlon and Messenger, 1998) with a mass die off after the breeding season. However, the Brittiany (north west France) population have a slightly different life history with two breeding groups. They are labelled as GIB (group one breeders with a shorter 12-15 months life cycle) and GIIB (group 2 breeders, a longer life cycle of 18-24 months; Boletzky 1983; Gauvit et al. 1997). Each group appears to produce the next generations alternate group, so GIB produce GIIB and vice versa. This results in two overlapping generations variable for size and extent of maturity that due to times of highly skewed operational sex ratios cannot be reproductively separated from one another (Gauvit et al., 1997). A comparable situation may exist in the Bay of Biscay, NW Spain (Guera and Castro, 1988), in *S. apama* in northern Spencer Gulf (Australia; Hall et al., 2007) and possibly in *Sepia latimanus* (Dan et al., 2012).

## 2. Materials and Methods

Video 1 (supplementary materials) was recorded from SCUBA dives off the south Cornish coast (UK) in early September 2013. Movie S1 comprises of a single male signalling to a single female, within a group of six. The entire dive video lasted approximately 55 mins, taken in late afternoon and comprises a group of six *S. officinalis.* The behaviour of interest lasts for approximately 2 minutes. The video was not recorded for data collection purposes, rather for public media use with the behaviour of interest only noticed later by the first author of this paper. The fortuitousness of obtaining the video prompted a citizen science request for more videos that may contain behaviours not published before. A request for videos of UK cuttlefish (i.e. *S. officinalis*) was posted on a UK based Facebook group “UK Viz Reports” (an active, closed Facebook group that shares visibility reports, general diving conditions and videos of UK marine animals daily with >6k members) received multiple replies. From these replies, a second video (Video 2) was obtained, again filmed just off the Cornish coast (UK), but this time in September 2017. Additionally, a third video (Video 3) was obtained from August 2017, again from roughly the same area of the Cornish coast. Videos were chosen for analysis based on there being groups (n>2 individuals) of sexually mature cuttlefish or that possessed examples of relevant behaviour. Stills were taken from the videos using Microsoft Office^TM^ software.

## 3. Results

### 3.1 Arm hiding (female mimicry) and ‘split signal’ (sneaker)

Video 1 appears to show a male attempting to deceive rival males (out of shot) by hiding two of its outer arms (R3 and R4) whilst courting a female – see figure 1 (C-F), Movie S1 (supplementary material. The video begins what appears to be a male (based on its subsequent behaviour) and a possible female. Three more cuttlefish can be observed to the left but do not always appear in shot, do not interact with the two focal cuttlefish and play no part in the analysis provided here. On close inspection, the y female has relatively slight damage to her arms (sucker marks) but no significant mantle damage which is common in this species towards the end of their breeding season (c.f. Allen et al., 2017, Video 2 this paper and figure 2F). It has the posture of a female about to spawn (see Hall and Hanlon 2002 figure 9 a, b for a similar posture in *S. apama*) or has just recently spawned. The two focal cuttlefish hover approximately 1 meter above the seafloor in more or less the same position throughout the two minutes of footage. After 34 seconds (0:34 on the video) the cuttlefish on the left begins to pull up arms R3 and R4 into the mantle cavity (figure 1C). This action occurs suddenly, and the entire process takes only a few seconds. During the behaviour the right-hand side of this cuttlefish appears to exhibit a weak zebra pattern, it is unclear which pattern is displayed on the left-hand side of its body. The leucophores and iridophores on the arms L3 and L4 remain visible and appear iridescent, at least to human eyes. On hiding its arms, the cuttlefish increases its mantle fin oscillations, potentially indicating heightened awareness or anticipation (Cooke and Tonkins 2015). Upon hiding arms R3 and R4, the cuttlefish appears to be leaning towards the second cuttlefish (figure 1F) from hiding its arms until releasing them again. At 1:14 arm L4 appears to be showing intense zebra patterning but unfortunately the rest of this side of the body is out of shot. The hiding of the arms ends at 1:27 and again happens suddenly with the arms reappearing quickly. The cuttlefish on the right appears unchanged throughout the video footage apart from small amounts of movement and occasional R1 and L1 arm twitching. No copulation takes place, nor does the behaviour appear again throughout the entire dive (~55 minutes). From the same footage a still (figure 1A) was taken showing what appears to be a split signal, as described in (Brown et al., 2012) which is similar to a photograph of the same species taken in an aquarium (figure 1B).

**Fig 2.**
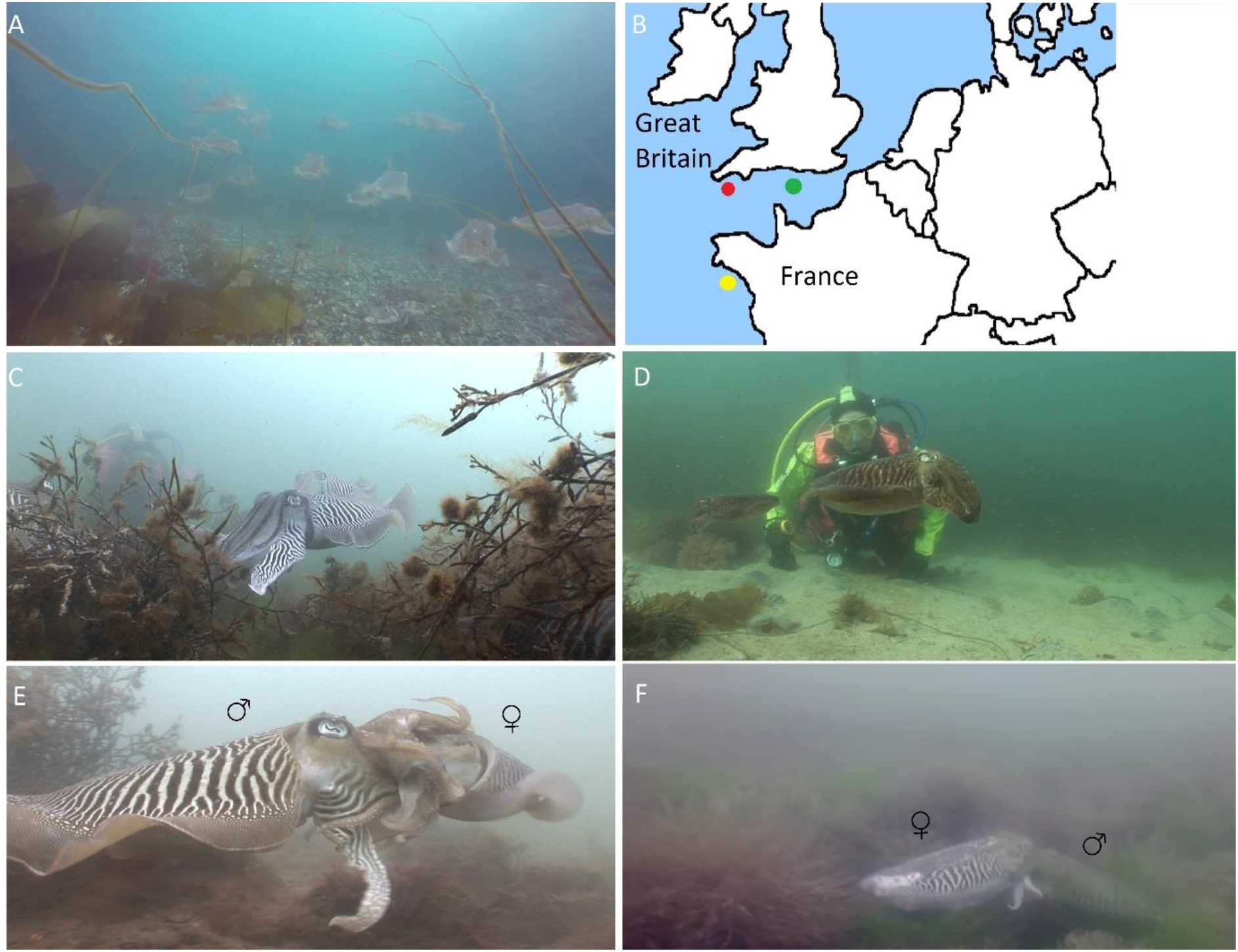
A) Video still of ~ 18 *S. officinalis* from a group of ~ 30 taken from a recording off coastal Cornwall, UK in September 2017 – see movie S3. B) locations of the cuttlefish studied here (red dot), biennial breeding group in the English Channel (green dot; Bloor et al. 2013a) and the long/short life cycle group (yellow dot; Gauvit et al. 1997). C) Still taken from video recorded in June 2013 showing large and conspicuous males c.f. D) Still of a male taken from video recorded at the beginning of September 2013 which also includes sneaky behaviour in fig 1 C-F and Movie S1. Note: same diver is caught in both stills and provides approximate size differences in the cuttlefish. E) Male (left) mating with female he has been mating guarding, cf. F) male (right) from Movie S2 who opportunistically mated with a female whilst two males competed for her

### 3.2 Spontaneous copulation video

Video 2 begins with two male cuttlefish (based on behaviours and constant use of intense zebra patterning) observed in competition for a female cuttlefish (below right of the males, her sex based on her smaller size, later copulation with a presumed male, lack of use of arms L1 or R1 in communicating with the other cuttlefish and significant mantle damage – see Allen et al., 2017). The two males are using intense zebra patterning, dark eyes (c.f figure 1 C-F, but see Allen et al., 2017) and waving their outer arms at one another in agonistic signalling. One of the males, closest to the female, appears to be mate guarding, which might suggest the pair had recently copulated. At 0:10 one of the males backs away and the pair move out of shot. The film now cuts to a scene like the one preceding it but with a fourth (presumably male) cuttlefish joining from the left – 0:28). This new cuttlefish is exhibiting the intense zebra patterning and darkened eyes. However, the intensity of the dark eyes seems to be less intense than that of the two other males engaged in antiparallel circling with 4^th^ arm extensions, typical agonistic displays for this species (see Allen et al., 2017; Hanlon and Messenger, 1998). At 0:34 the new male, still exhibiting the intense male patterning, ducks below the two males and forces copulation with the female. His patterning changes dramatically from intense zebra patterning to mottle (0:36), then flushes dark brown (0:37). The copulating male and female circle briefly, below the two still competing males, before moving to the sea floor with the male exhibiting weak zebra patterning (figure 2F but c.f. figure 2E) until the video ends at 0:46.

### 3.3. Large group of cuttlefish

Video 3 (and figure 2A) provide evidence of large groups of mature cuttlefish in UK waters. The video footage begins with what appears to be nine cuttlefish of a similar size, in a line, floating approximately 2-3m off the sea floor. This formation is not unusual in groups of *S. officinalis* (Cooke pers.obs) in the wild but its function is unknown. The formations seen in the video closely match that seen in *S. latimanus* (Yasumuro, et al., 2015). Within this sub group there are minor interactions resembling mild warning displays (deimatic eye spots) when one cuttlefish gets too close to another. The group of nine are slowly moving right (from the cameras point of view), looking like they are headed somewhere. The camera briefly follows a lone individual (possibly female) before panning right, and from 1:25 many more cuttlefish are revealed. This time the cuttlefish are closer (~0.5m) to the sea floor but go up as far as 3-4m in the water column, again resembling observations seen in *S. latimanus* (Yasumuro, et al., 2015). Although the video footage is quite poor a minimum of 15 cuttlefish can be made out between 1:25 and 1:30. Again, interactions are minimal, minor warning displays (deimatic eye spots for e.g.). It is not possible to definitively sex many, if any, individuals but they all appear to be sexually mature based on a) some zebra patterning (see 1:47) b) swimming behaviour (i.e. above the sea floor) and c) the relative size of the cuttlefish. At 1:49 it is possible to count 19 individuals (again with some in shot exhibiting zebra patterning). No feeding is observed, nor any courtship/mating. The camera person appears to alter the behaviour of individuals by getting too close, but generally they appear to be moving in one direction, consistently and as a group (again see *S. latimanus* (Yasumuro, et al., 2015)). The video ends at 2 mins. The divers who recorded the film were able to count 30 individuals, with there likely being more were too difficult to count in the circumstances.

## 4.0 Discussion

We believe the results presented in this paper are potentially very interesting and the observations promote much further discussion. Figure 1 (A and B) seems to show the split signal, as reported for *S. plangon* (Brown et al. 2012) in the wild and in the aquarium for *S. officinalis*, the first time this has been reported in this very well studied species. Furthermore, Figure 1 (C-F) and Video 1 suggests *S. officinalis* may use the arm hiding deceptive behaviour reported in *S. apama* (Hanlon et al. 2005; Norman et al. 1999), again a first for this well-known cephalopod species. Although no copulation takes place, and the female looks uninterested, alternative explanations for the behaviours are not immediately forthcoming. Given the relative size of the animals (relatively small), the time of year (September), it is perhaps plausible that the cuttlefish are GIIB long cycle individuals. This would mean they have just reached sexual maturity, or on the cusp. The lack of success, or even acknowledgement, of the behaviour by the female may indicate ‘she’ is not fully mature yet – females do typically mature a little later (Hanlon and Messenger, 1998). Many animal species play at copulating behaviours (e.g. Bonobo chimpanzees (*Pan paniscus*) Kuroda 1980) and the behaviour seen here might be considered play, in the sense play is a form of practice. The female may have the posture of just spawned, or about to spawn (see Hall and Hanlon 2002 figure 9 a, b for a similar posture in *S. apama*) which may also explain her disinterest. The unresponsive female does not have the severe mantle damage, as seen frequently in very old female cuttlefish (e.g. Allen et al., 2017), but sucker marks can be seen on her head, suggesting a male has tried to copulate with her at some point.

Video 2/Fig 2F shows a male opportunistically mating a female whilst two other males compete for her. On grabbing hold her, he adopts a brown body colour, this is different to copulations when males normally maintain the intense zebra pattern (figure 2E) followed by mate guarding and a change from his arrival during which he was signalling the intense zebra pattern. The schematics (figure 3) attempt to neatly represent the nature of *S. officinalis* sneaky mating when Movie S2 shows the opportunistic nature of males. In this video, the male is comparable in size to the other males and uses conspicuous aggressive colouration right up until copulation, when he quickly reduces conspicuousness (i.e. does not exaggerate his features and reduces the conspicuousness of his patterning). The behaviour might be considered sneaky (i.e. and AMT) in that the male uses the competing males for cover and then reduces its conspicuously male traits (intense zebra pattern) during copulation (c.f. figure 2E and Allen et al., 2017 with figure 2F).

**Fig 3.**
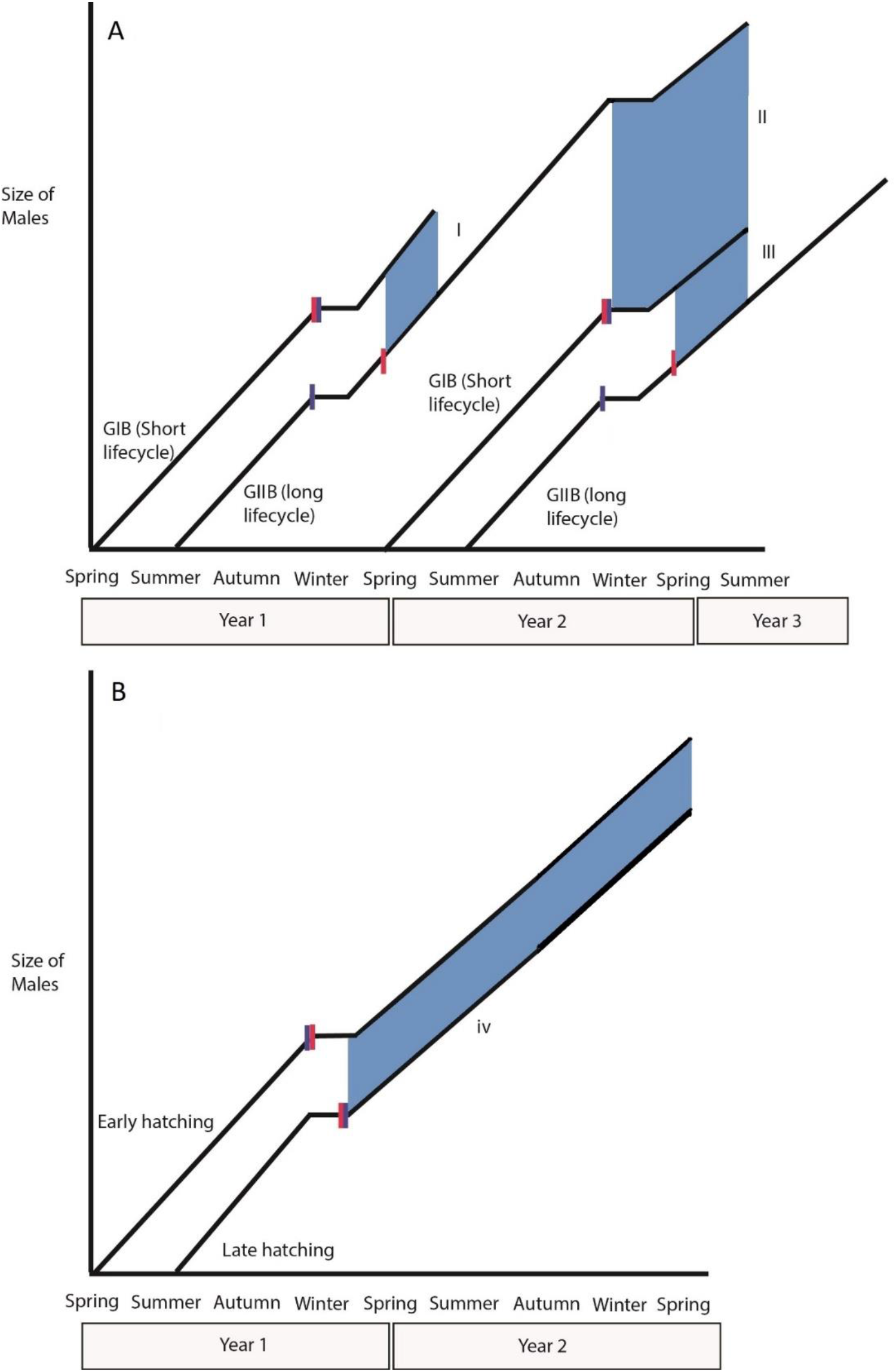
Schematic of two proposed lifecycles in S. *officinalis* based on information from Gauvit et al. (1997; NW France), Guera and Castro (1988; NW Spain) and Bloor et al. (2013; English Channel). A) Two breeding groups (Gauvit et al. 1997) scenario which are geographically proximate (Brittany, NW France, see figure 2B) to the south Cornish (SW England) population studied here. A) Two groups named GIB (group one breeders) and GIIB (group two breeders). In this schematic GIB Breeders coincide (blue shaded area) with small GIIB Breeders (I) when both are sexually mature (blue shaded zones). B) Alternatively, the population may be biennial, based on their proximity to the English Channel (Bloor et al. 2013a; see figure 2B). This lifecycle might encourage sneaky mating in males hatched later in the year and/or who were less successful in obtaining food. Blue shaded area = approximate duration of overlapping mature males. Red bars = approximate time of sexual maturity (Boletzky 1983; Dunn 1999) and blue bars = body growth stops over winter (Dunn, 1999) but gonad growth and sexual maturity continue, some mating’s do take place over winter (Guera and Castro 1988).

*S. officinalis* have previously been described as solitary or comprising of small (n ~ six) groups (Hanlon and Messenger 1998). Figure 2A, Video 3 shows groups (n ~ 21/30) that could further drive AMTs if competition for mates is intense. However, larger groups may hinder the split signal/arm hiding strategy which requires the rival cuttlefish to be in specific places relative to the signaller (Brown et al. 2012). Large groups may encourage more frequent alternative strategies such as males adopting a whole-body female like phenotype as seen in *S. apama* (e.g. Hanlon et al. 2005; Norman et al. 1999). The video evidence suggests it is possible that *S. officinalis* populations may aggregate/lek at unknown and/or transient locations similar in nature to closely related species (e.g. *S. apama* Norman et al. 1999) but await discovery. The grouping could be considered a school (see Krause et al., (2000) for definition of what a ‘school’ might be), as seen in *S. latimanus* (Yasumuro, et al., 2015), rather than an aggregation for breeding per se. The observation that the cuttlefish are schooling would also be a first for this species and deserves further investigation.

### 4.1 Conditions for alternative mating tactics

Figure 3 shows two schematics based on possible life histories of the cuttlefish seen here (see Gauvit et al. 1997 and Bloor et al. 2013a). GIB Breeders coincide with smaller GIIB Breeders in Spring/early Summer when both are sexually mature (Figure 3Ai). GIIB males, the next year and much larger, again meet new GIB males who now are smaller (Figure 3Aii/iii). This provides GIIB males conditional strategy opportunities: when small they could adopt the female mimic strategy, but when larger than GIB males they can adopt the normal dominant status strategy. The same is true for GIB males who may find themselves with larger or smaller GIIB males. Alternatively, if the Cornish population are like the English Channel biennial breeders (Bloor et al. 2013a) individuals will still come across males both larger and smaller, unless they are at the extreme end of the size spectrum (i.e. hatched very early or very late). The alternate life cycles are not unique to *Sepia officinalis* at this location, they are found in the Bay of Biscay, NW Spain (Guera and Castro, 1988), in *S. apama* in northern Spencer Gulf (Australia; Hall et al., 2007) and *S. latimanus* (Dan et al., 2012).

It is unclear how OSRs may influence AMTs in cuttlefish. Hall and Hanlon (2002) reported an extreme OSR in *Sepia apama* of 11:1 (m:f with an average of 4:1) yet some species have been reported to have female skewed (*Sepia pharaonis* = 1:3 (Abdussamad et al. 2004; but c.f Silas et al 1986)). OSRs in cuttlefish can vary dramatically within a species. Rao (et al., 1993) reported a very wide range of OSRs (33:67 – 70:30; m:f) in *Sepia aculeate*, suggesting that AMTs may not be present at all in some populations of the same species if OSR is a major factor in their use.

In lieu of any genetic evidence, and using the data that does exists regarding their biogeography/ecology/life cycles (i.e. Gauvit et al. 1997), and possibly variable OSRs, the decision to express an AMT is probably conditional based, rather than inherited. This is like AMTs in guppies (Reynolds et al. 1993; or the dung beetle *Onthophagus* sp., Thornhill, 1981), which are environmentally driven ‘conditional’ strategy (Gross 1996; Neff and Svensson 2013. This differs to genetically determined ‘pure’ strategy (e.g. the sword tail *Xiphiphoorus nigrensis,* Ryan et al 1992; the ruff *Philomachus pugnax*, Lank et al. 1995). The isopod, *Paracerceis sculpta,* has three male alternative strategies; alphas guard territories and females; betas adopt a female phenotype; gammas sneak (Shuster 1989). The strategies are genetically determined (1 gene, 3 alleles; Shuster 1989) but similarly describes the three strategies seen in *Sepia spp.* Coleoid cephalopods are unique as to the extent at which they can alter their appearance, being able to adopt most possible phenotypes, almost instantaneously and this possibly removes the requirement of fixed phenotypes determined genetically.

### 4.2. Tactical deception and eavesdroppers

Hiding secondary sexual characteristics in the way the cuttlefish appear to do here and in *S. apama* (Hanlon and Hall 2002) are used to deceive males who are possibly eavesdropping. A deceived male cuttlefish would pay a high cost of being unwittingly usurped (see Norman et al. (1999) for relatively high fertilization rates of sneakers in *S. apama*). The consequences for the sneaker might be considerable if caught (although fights rarely escalate to physical confrontation, Allen et al., 2017), and then injured or worse, in an ensuing fight, suggesting this would be genuine form of deception (Kuczai et al. 2001). de Wall (1998) found that a male chimpanzee (*Pan troglodytes*) would hide his erection when dominant males were in view, whilst simultaneously allowing a female to see it (de Waal, 1998), a situation described as tactical deception with claims the animal must possess a ‘theory of mind’ (Hare 2000; Kirkpatrick 2007, and already being stated as present in a cuttlefish – Cuthill 2007). If initially smaller males do transition into larger, dominant and conspicuous males, then sneakers who use deception may remain at a low enough frequency for the honest signals (intense zebra patterning/larger size) to remain a useful means in determining male dominance (Johnstone and Grafen 1993).

Cuttlefish are polyandrous (Naud et al. 2005), and females in this genus appear to accept sneaker males for egg fertilisation (Norman et al. 1999) and may even prefer males who use non-threating zebra patterning. Boal (1997) found females were repelled by males exhibiting the intense zebra patterning; they were possibly wary of receiving aggression, which is presumably how males interpret the signal (Boal, 1997). Sexually selected honest signals in male competition could be helping drive an alternative phenotype consisting of a less aggressive signal that is more appealing to females. When the split signal is employed (Brown et al. 2012; Figure 1A/B) it is notable that the signal on the side of the female is not as intense as the signal used in male-male competition (c.f figure 2E, Allen et al., 2017).

### 4.3 Are AMTs common in cuttlefish?

Many other *Sepia* species (over 100 have been described, see Reid et al. 2005) have similar life history properties that may encourage selection for alternative mating strategies, such as; over lapping generations where smaller males may be competing for mates with bigger males; inshore migrations that increase densities of mature animals; conspicuous male courtship signals/sexual dimorphism and mate guarding; limited spawning sites; all of which provides the opportunity for alternative male phenotypes – see table 2. Given the commonality of these key features we predict that alternative strategies, such as sneaky copulations via tactical deception and or female mimicry, may be more common in this genus than thought. Exaggerated secondary sexual traits can drive AMTs via sexual selection (Neff and Svensson 2013), and these traits may be a sign post as to which species should be further investigated (table 2). Many *Sepia* species, not previously reported in this context, possess the ability to produce conspicuous displays, even by cuttlefish standards (e.g. *Sepia latimanus*, *Sepia prashadi*, *Sepia mestus*, *Sepia omani*, *Sepia filibrachia* etc). There is no evidence that these hyper conspicuous displays are aposematic warning signals and their use in predation is unclear, so they could at least in part be driven by sexual selection and deserve further attention.

**Table 2:**
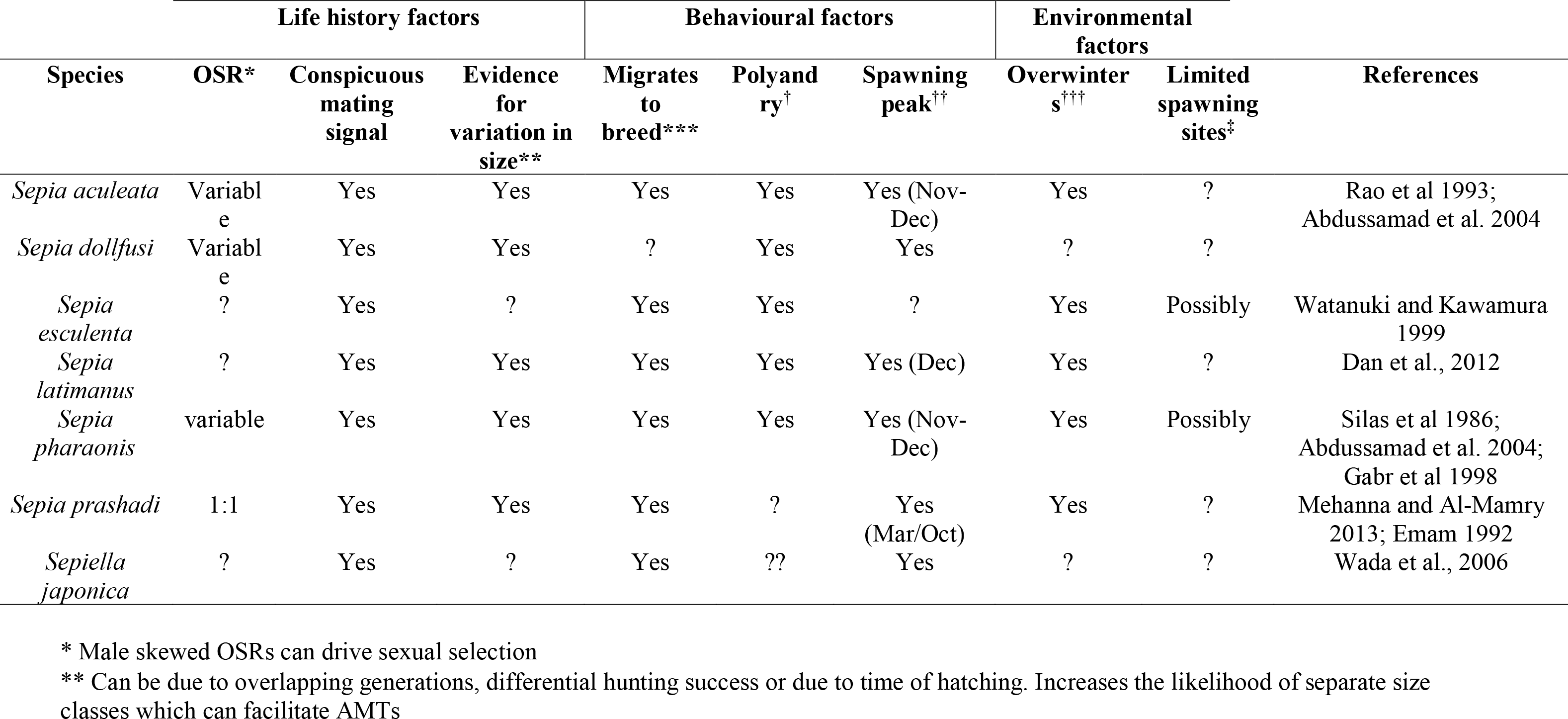
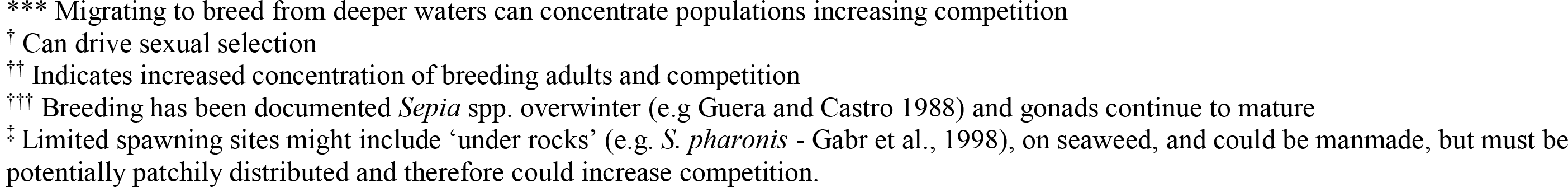
Information, where available, pertaining to the likelihood of AMTs being present in other *Sepia* species. Includes information on a closely related genus (*Sepiella*) as they may also prove to exhibit AMTs. Does not include *S. officinalis, S. apama* nor *S. plangon* for brevity, nor most of the remaining 100 species where nearly all relevant data is missing. Factors are based on life history, behaviour and the environment, all of which could drive sexual selection and ultimately the evolution of AMTs. Data collected almost exclusively through fisheries studies, highlighting their usefulness when studying *Sepia* spp. in the context of evolutionary biology. However, given the wide biogeographic ranges of many coastal species, fisheries data is only useful for the location it describes

| Species | Life history factors |  |  | Behavioural factors |  |  | Environmental factors |  | References |
| --- | --- | --- | --- | --- | --- | --- | --- | --- | --- |
|  | OSR* | Conspicuous mating signal | Evidence for variation in size** | Migrates to breed*** | Polyandry<sup>†</sup> | Spawning peak<sup>††</sup> | Overwinter s<sup>†††</sup> | Limited spawning sites<sup>‡</sup> |  |
| *Sepia aculeata* | Variable | Yes | Yes | Yes | Yes | Yes (Nov-Dec) | Yes | ? | Rao et al 1993; Abdussamad et al. 2004 |
| *Sepia dollfusi* | Variable | Yes | Yes | ? | Yes | Yes | ? | ? |  |
| *Sepia esculenta* | ? | Yes | ? | Yes | Yes | ? | Yes | Possibly | Watanuki and Kawamura 1999 |
| *Sepia latimanus* | ? | Yes | Yes | Yes | Yes | Yes (Dec) | Yes | ? | Dan et al., 2012 |
| *Sepia pharaonis* | variable | Yes | Yes | Yes | Yes | Yes (Nov-Dec) | Yes | Possibly | Silas et al 1986; Abdussamad et al. 2004; Gabr et al 1998 |
| *Sepia prashadi* | 1:1 | Yes | Yes | Yes | ? | Yes (Mar/Oct) | Yes | ? | Mehanna and Al-Mamry 2013; Emam 1992 |
| *Sepiella japonica* | ? | Yes | ? | Yes | ?? | Yes | ? | ? | Wada et al., 2006 |

\* Male skewed OSRs can drive sexual selection
\*\* Can be due to overlapping generations, differential hunting success or due to time of hatching. Increases the likelihood of separate size classes which can facilitate AMTs
\*\*\* Migrating to breed from deeper waters can concentrate populations increasing competition
† Can drive sexual selection
†† Indicates increased concentration of breeding adults and competition
††† Breeding has been documented *Sepia* spp. overwinter (e.g. Guera and Castro 1988) and gonads continue to mature
‡ Limited spawning sites might include ‘under rocks’ (e.g. *S. pharonis* - Gabr et al., 1998), on seaweed, and could be manmade, but must be potentially patchily distributed and therefore could increase competition.

If these predictions prove correct, *Sepia* spp. could make an excellent model for the study of alternative reproductive strategies. Not only would there be many accessible natural replicates (technology now exists to tag and or track cuttlefish in the wild; Bloor et al. 2013b, Dan et al., 2012), but the techniques for rearing and experimenting with cuttlefish in captivity are very well known. Additionally, with *S. officinalis* genome sequencing in progress, we may soon have the tools to learn more about the genetic nature of alternative reproductive strategies and which regions of the genome are under strong selection pressures in this group of animals, which is essential to further our understanding of sexual selection (Neff and Svensson 2013). Laboratory based reproductive studies involving *S. officinalis* invariably use single generations (e.g. Forsythe et al. 1994; Adamo et al. 2000; cf. brief mention of overlapping laboratory generations in Gauvit et al. 1997 p3) and roughly fed the same foods at the same amount, resulting in a single size classes (approximately, some individuals may remain very small – Cooke pers.obs). For many experiments having a single size class may be essential for replications so it is therefore not surprising that the behaviours reported here have not been seen before despite *S. officinalis* being extensively observed in captivity. Artificially creating distinct size classes may be relatively straight forward (see Boletzky 1983) and could be used to test the various hypotheses suggested here.

Some *Sepia* spp. have very wide ranges (e.g. *S. latimanus* ranges from Indo-West Pacific to southern Mozambique to Japan). They may even have had their reproductive behaviour observed (e.g. *S. latinmanus,* Corner and Moor 1980) but fisheries data (allowing us to infer the likelihood of AMTs) is often very specific to a relatively small area and should not necessarily be assumed to be true elsewhere for the same species (similarly, very little, if any, gene flow exists between allopatric populations *S. officinalis,* Perez-Losada 2002).

### 4.5 Use of citizen science

Citizen science is influential and widely discussed topic, has multiple origins and broadly refers to public participation and engagement in scientific projects (Riesch and Potter, 2014). Typically, projects are devised by scientists specifically with public engagement in mind with many important aspects (such as data quality and integrity) being potentially sacrificed or watered down to obtain the largest useful data set possible (Riesch and Potter, 2014). This might mean methods are simplified, accuracy removed or restrictions that allow untrained non-specialists to gather data. Through initial good fortune (video S1being shared with the lead author) we did not use a pre-designed data gathering method. Rather, we retrospectively requested data (via videos) which may remove some of the former issues (e.g. data integrity; Resnik et al., 2015). There are many benefits to using videos already collected by the public, including; access to a collection of footage from numerous locations uploaded free of charge to publicly accessible mediums (i.e. social media on the internet). In addition to specific benefits of our approach, broader benefits of citizen science remain, such as; meaningful engagement with the people who contribute to scientific funding (i.e. tax payers in the case of the UK); increasing the impact of science by making participants aware of local phenomena, understanding aspects of the scientific method and previously esoteric branches of science (Riesch and Potter, 2014). The downsides to our approach include; undirected random sampling; completely uncontrolled encounters with the animals and the data collectors interfering with the subjects; no standardisation of essential parameters; abrupt endings of footage and unhelpful angles when recording the animals. However, our approach does mitigate against an important general ethical issue with citizen science i.e. exploitation. All participants volunteered their videos and were happy to hand over intellectual property should it be an issue at the publishing stage. When used sparingly and with great care, working with public on wild animal observations can be of benefit to both parties. This paper would not exist without the videos collected by ‘amateurs’ and the participants (including other members of the Facebook group) learned a lot about a hidden and previously little understood species, and appeared to enjoy the experience. It’s telling that no quantitative analysis was possible and perhaps the most useful benefit of this approach to citizen science has been the ability to form hypotheses (e.g. that the *Sepia* genus may be a rich testing ground for sexual selection research) from videos of compelling behaviour. Should scientists wish to use our approach we recommend that clear, concise requests are made, that the publics knowledge is treated with respect, especially with respect to unusual observations that may at first seem unlikely.

## 5.0 Conclusions

*S. officinalis* appear to use signals and behaviours (split signal plus arm hiding), possibly attempting to deceive an eavesdropper and signalling to a potential mate. It appears AMTs may be more common in *Sepia* spp. than previously thought. This genus may have many ideal traits (accessible and predicted natural replicates; ease of lab rearing including short generation times; sexual dimorphism/conspicuous secondary sexual characteristics; fisheries data for many species; genome data soon available) for studying alternative reproductive strategies and sexual selection. ‘Citizen science’ has been shown to provide valuable observations of animals normally rarely seen, and there appears to be a vast amount of this resource on social media, populated by willing amateurs who are seemingly happy to engage with scientists.

## 6.0 Acknowledgements

The authors are grateful to the UK Facebook group ‘UK Viz Reports’ for their generous sharing of time and video footage. We are also very grateful for the two earlier reviewer’s comments who helped shape this manuscript with very useful suggestions and ideas. This work benefited from networking activities carried out under the COST ACTION FA1301, and is considered a contribution to the COST (European Cooperation on Science and Technology) Action FA1301 “A network for improvement of cephalopod welfare and husbandry in research, aquaculture and fisheries” (http://www.cephsinaction.org/).

